## Supplementary Figures for "Neuroblastoma signalling models unveil combination therapies targeting feedback-mediated resistance"

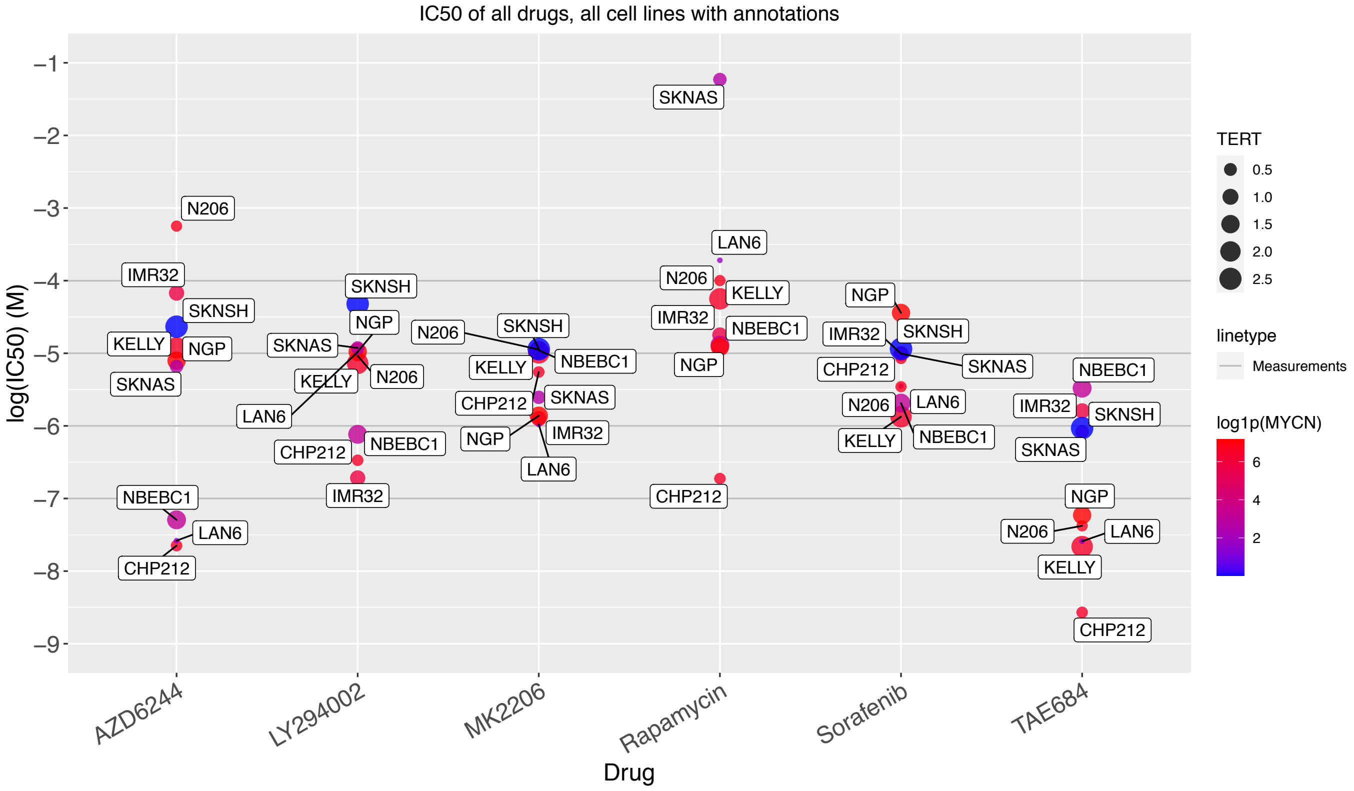


Supplementary Figure 1: Annotated IC50 of all measured drugs.

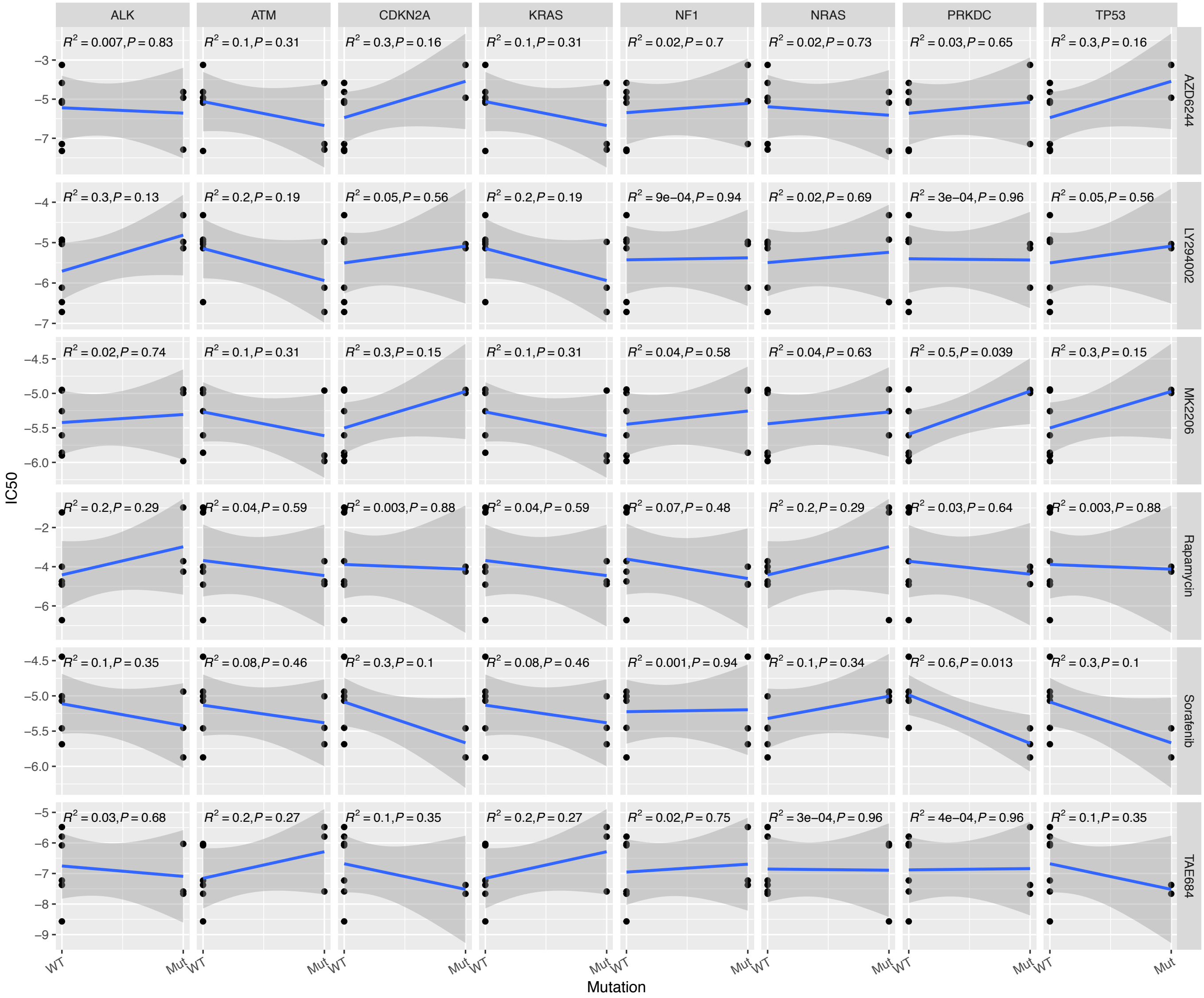


Supplementary Figure 2: (Correlation between IC50 and mutation status for RAS/P53 genes with mutation frequency between 30% and 70% in our panel.

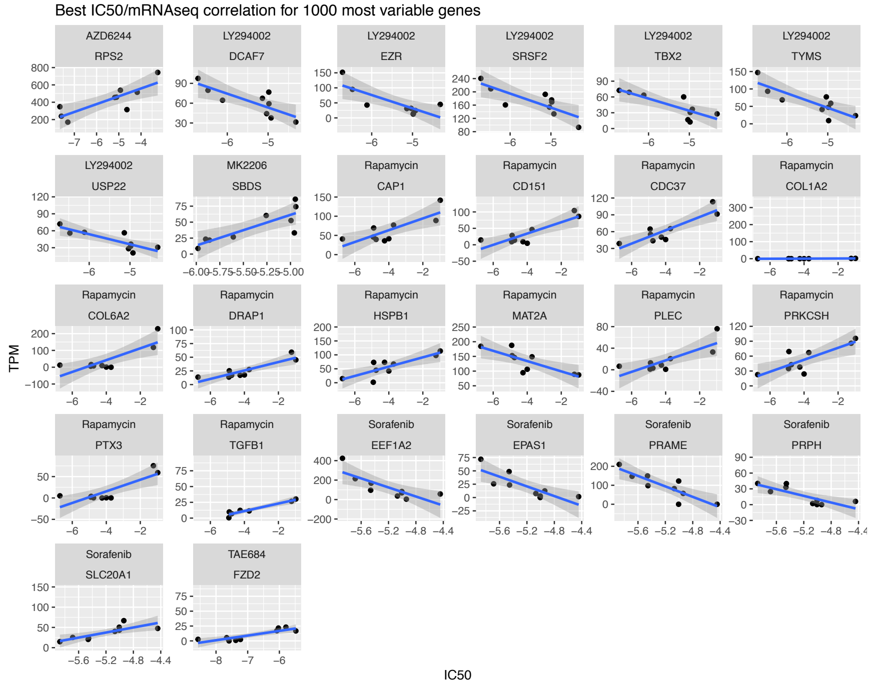


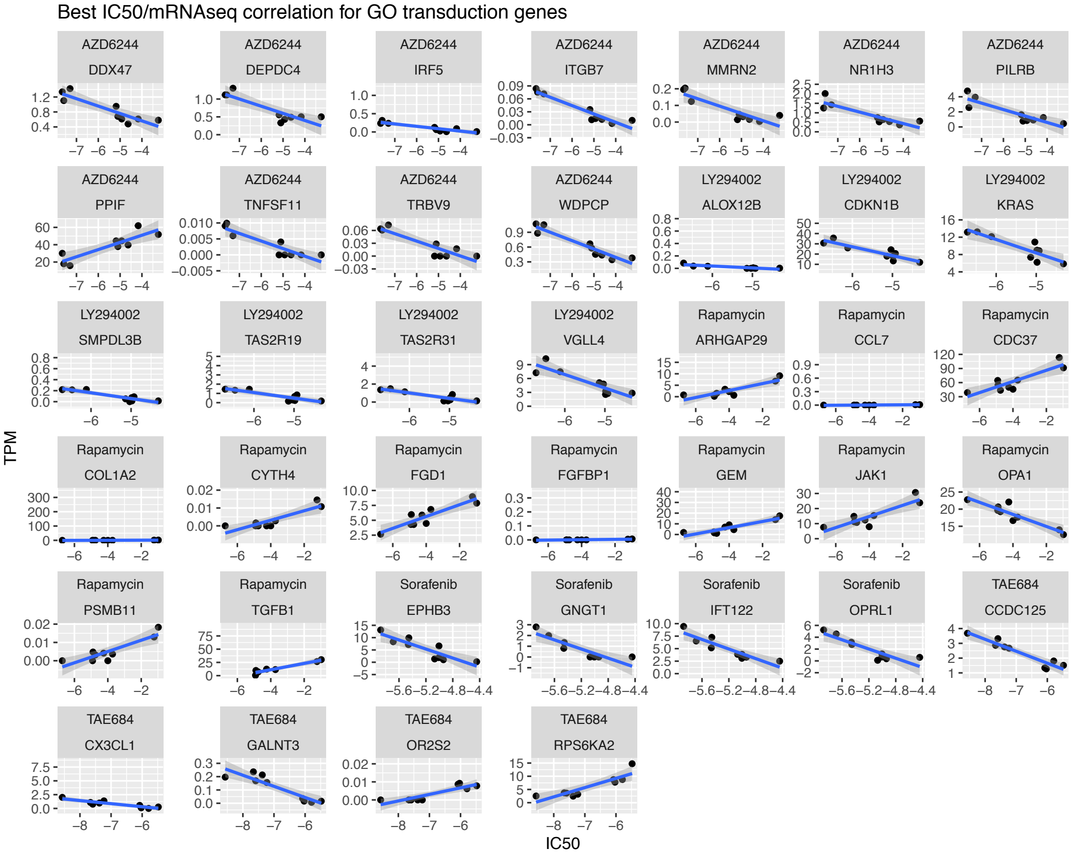


Supplementary Figure 3: Top correlations between IC50 and mRNA expression for the 1000 most variable genes (top, adjusted p>0.95) and GO signal transduction genes (bottom, adjusted p>0.95).

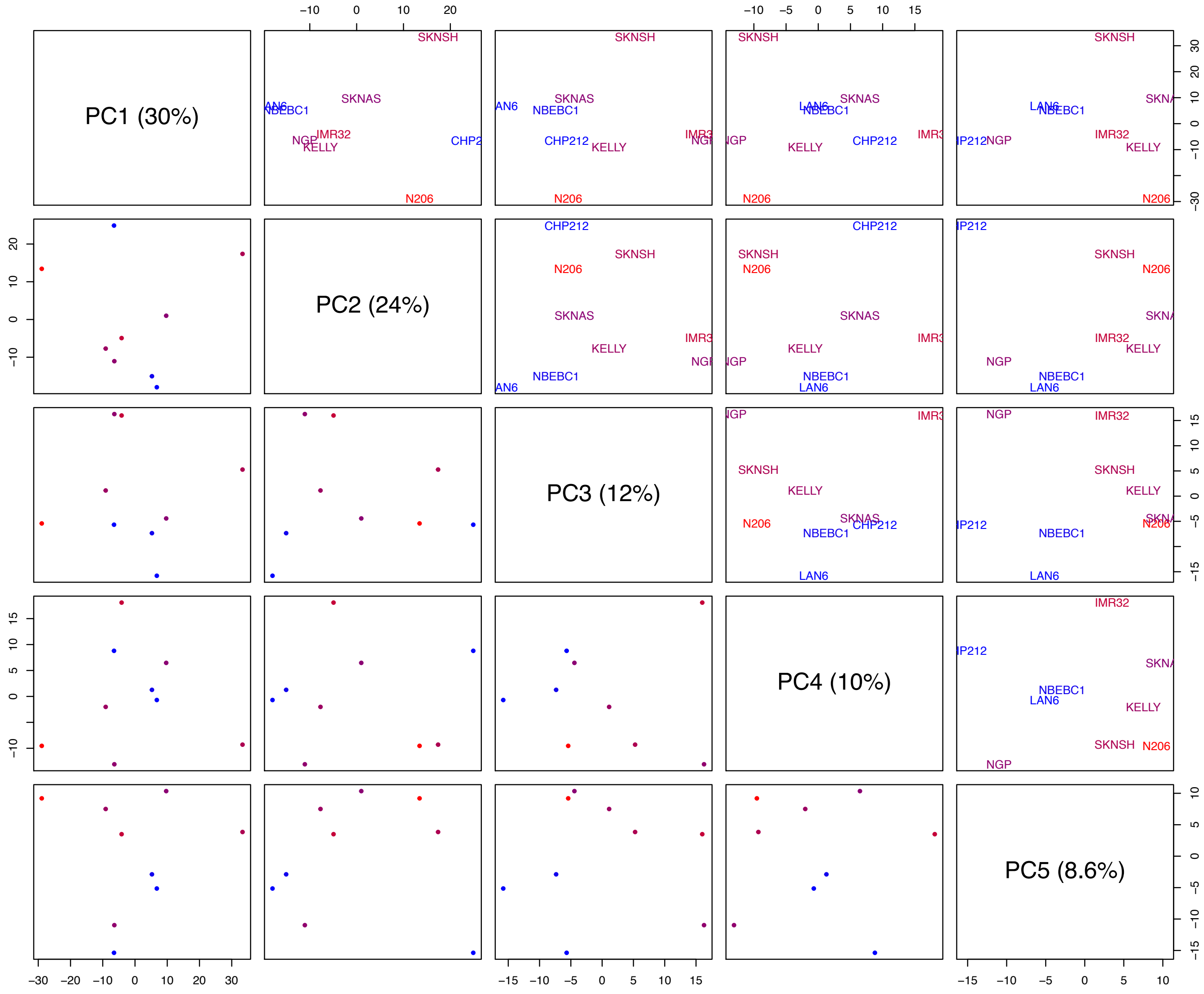


Supplementary Figure 4: Principal Component Analysis of the 1000 most variable genes. All components up to the first one explaining less than 10% of the variance are shown.

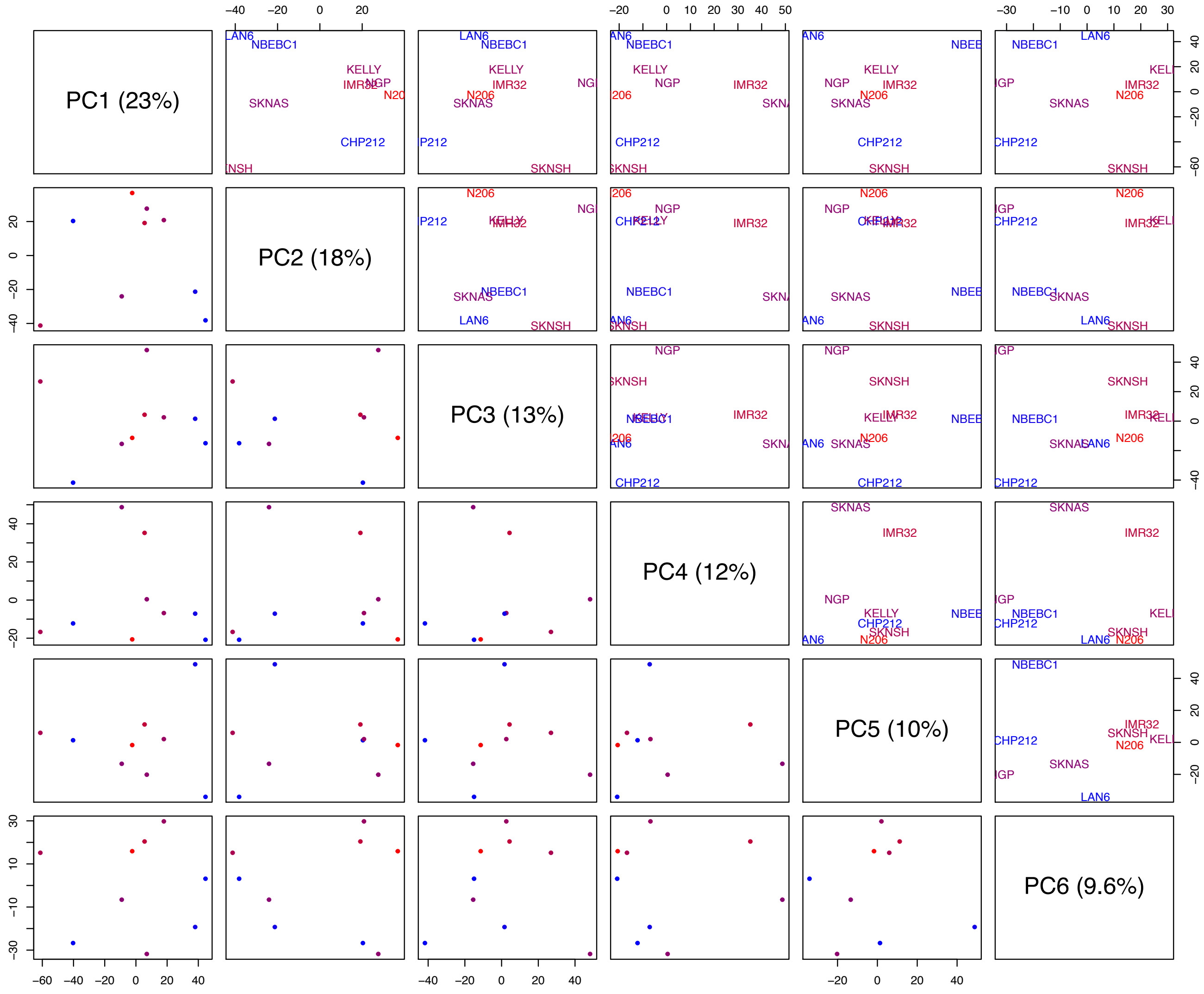


Supplementary Figure 5: Principal Component Analysis of the 5262 signal transduction genes as annotated in the Gene Ontology knowledgebase. All components up to the first one explaining less than 10% of the variance are shown.

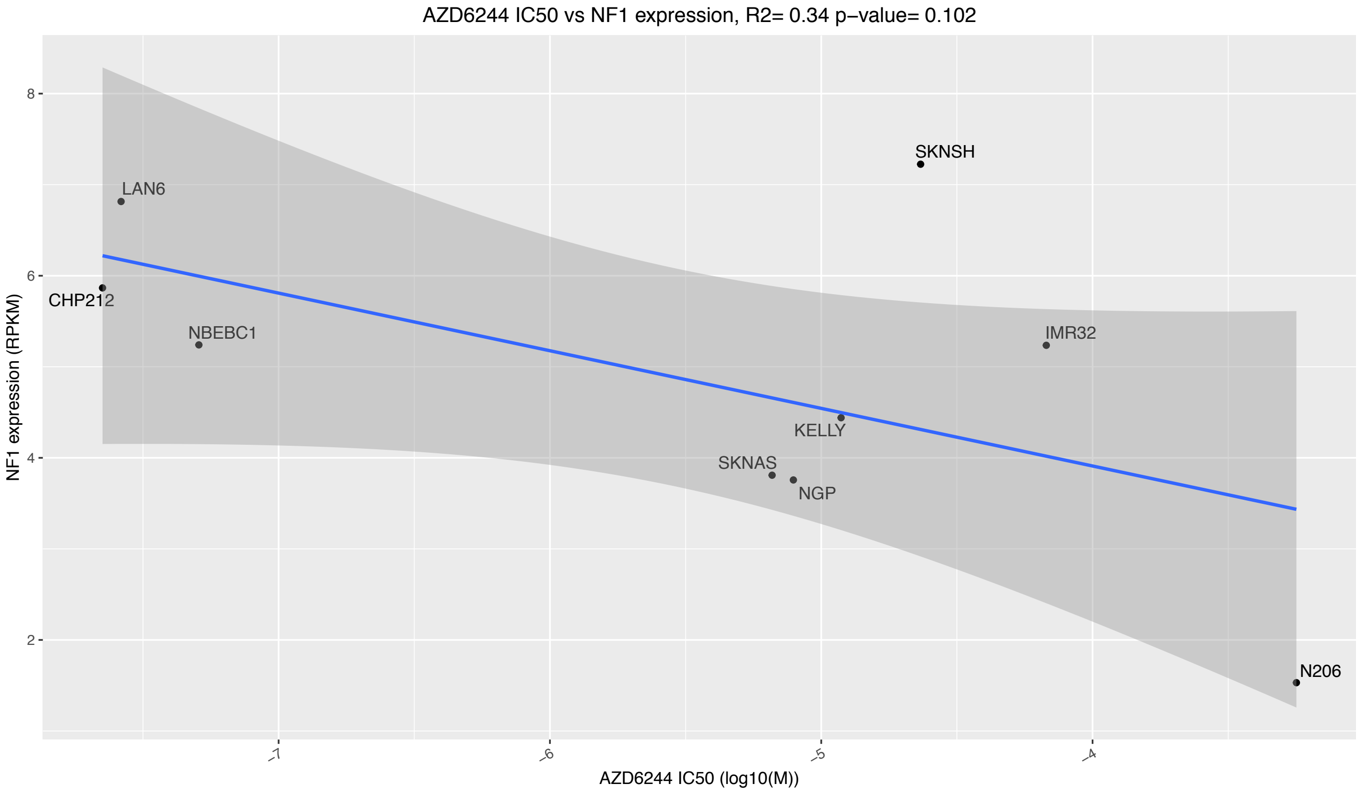


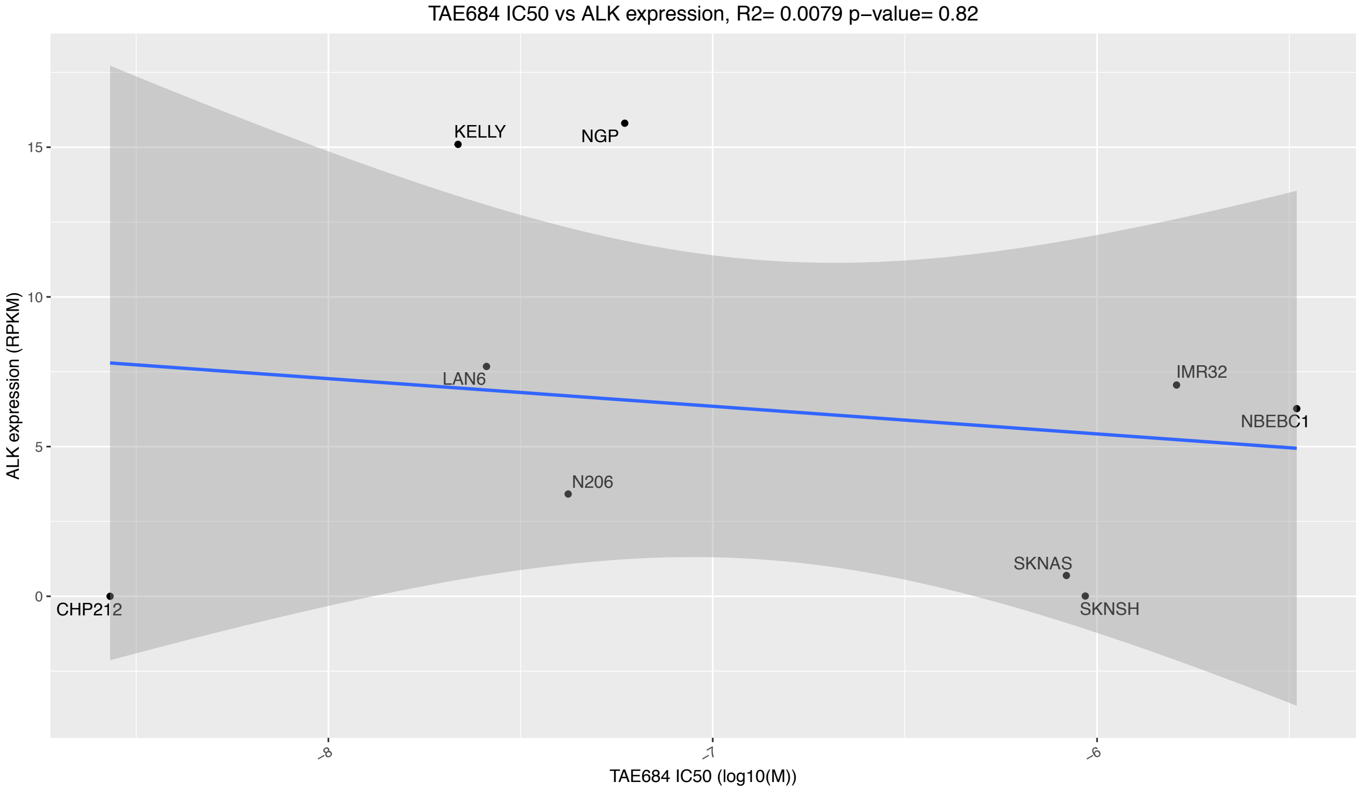


Supplementary Figure 6: NF1 expression versus AZD6244 IC50 and ALK expression versus TAE684 IC50.

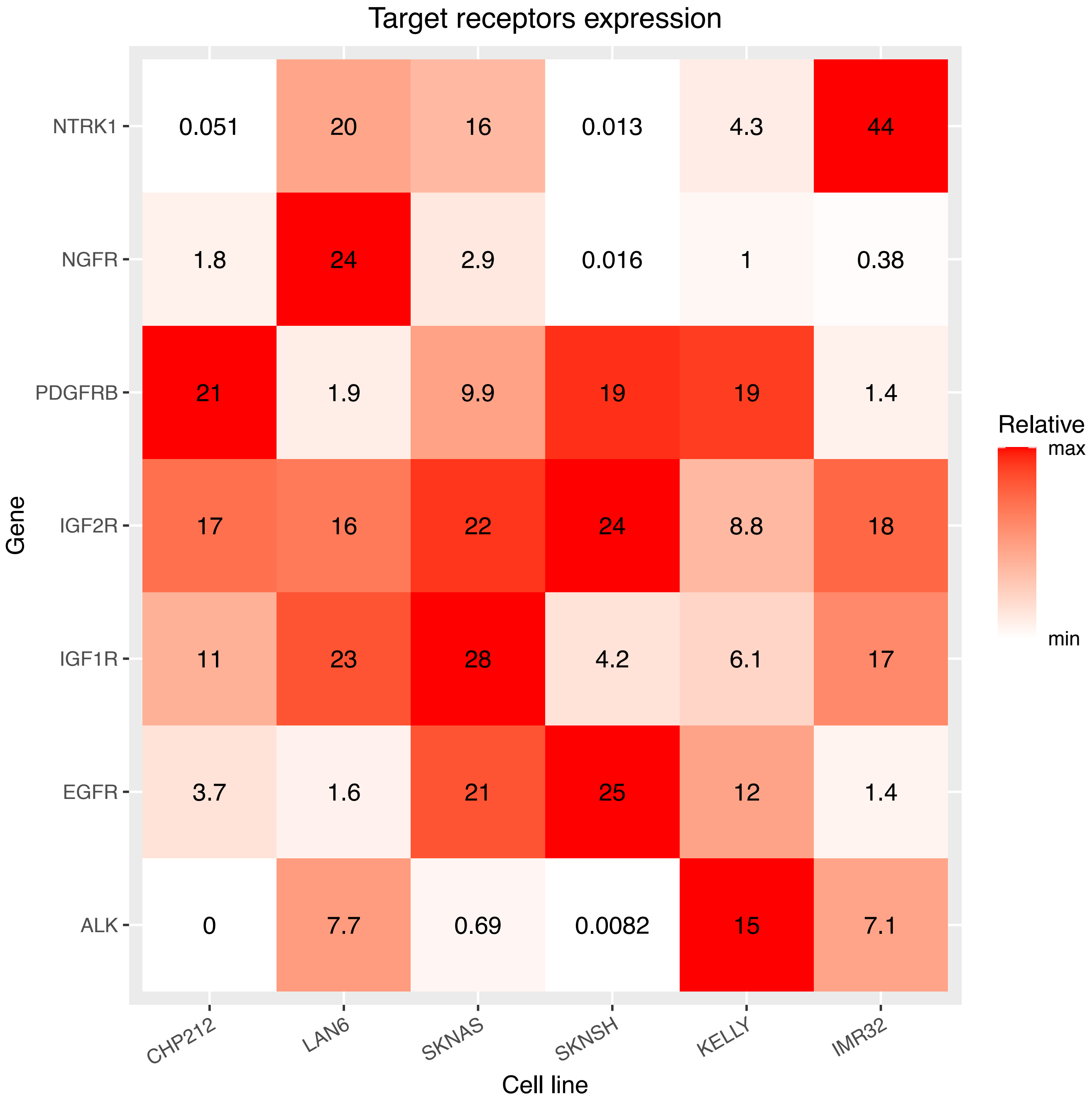


Supplementary Figure 7: Receptors RNA expression (TPM).

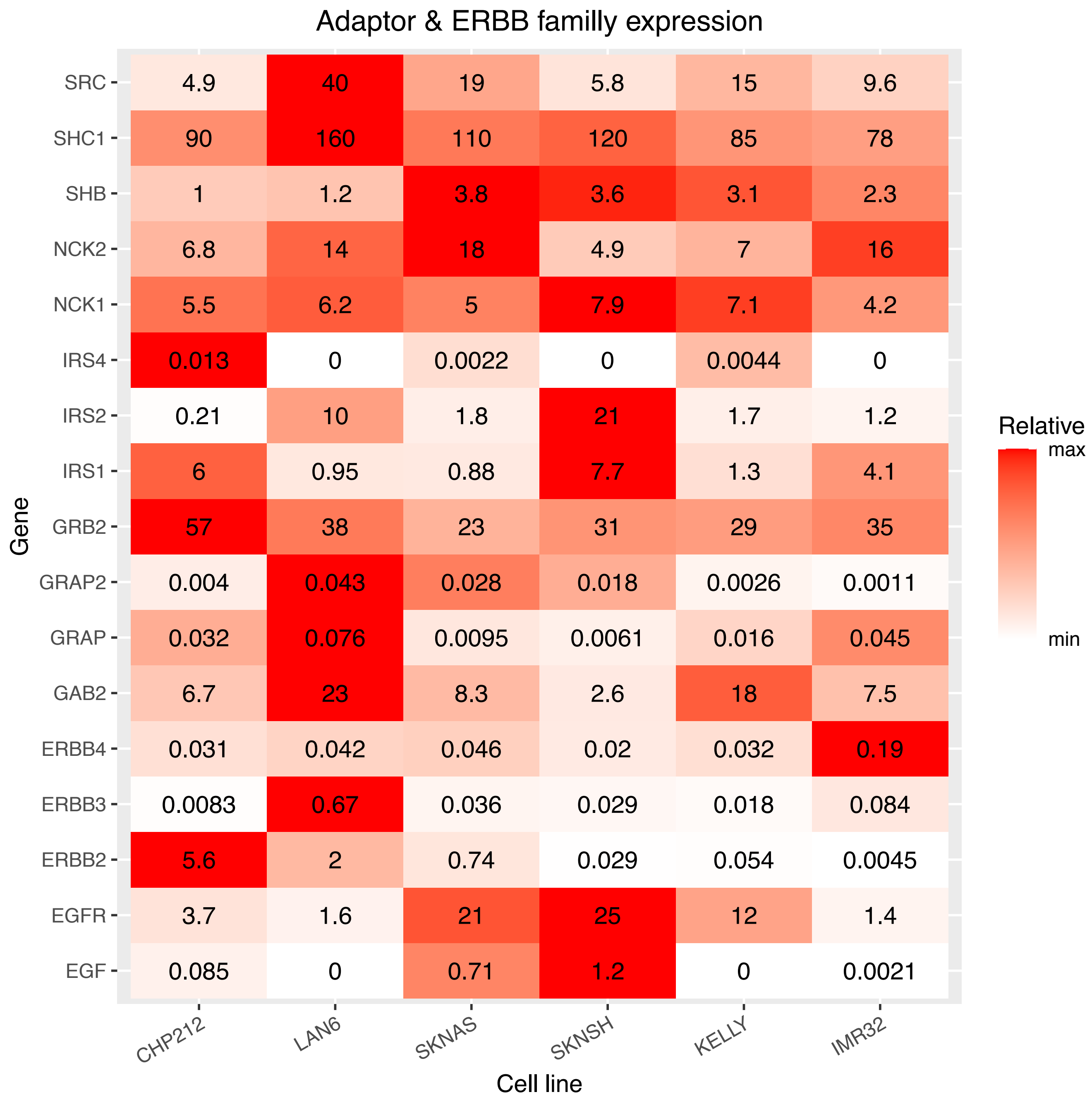


Supplementary Figure 8: Adaptors and ERRB receptor family RNA expression (TPM).

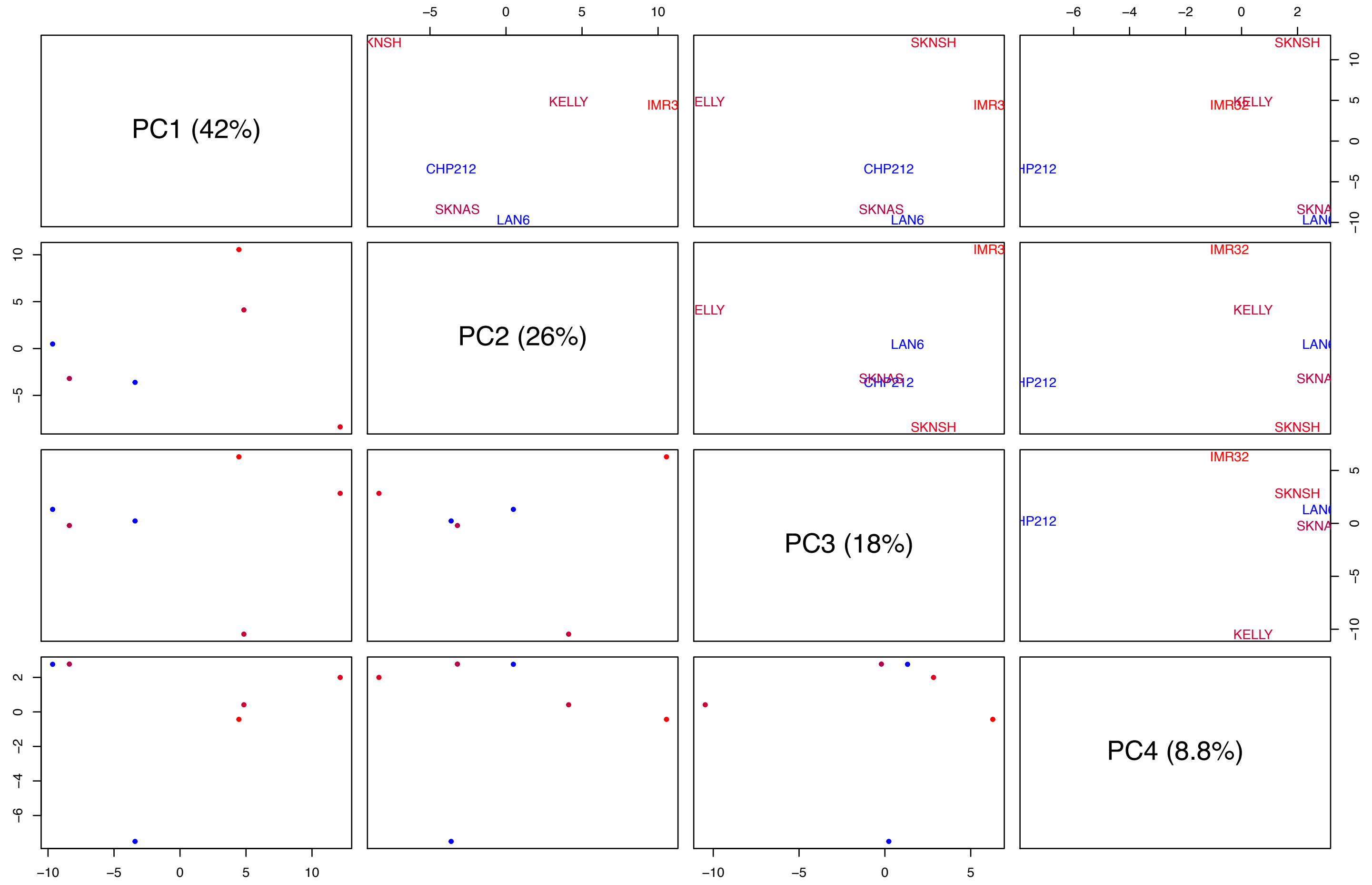


Supplementary Figure 9: Pair-plot of the principal components from the perturbation data in main Fig 2. All components up to the first one explaining less than 10% of the variance are shown.

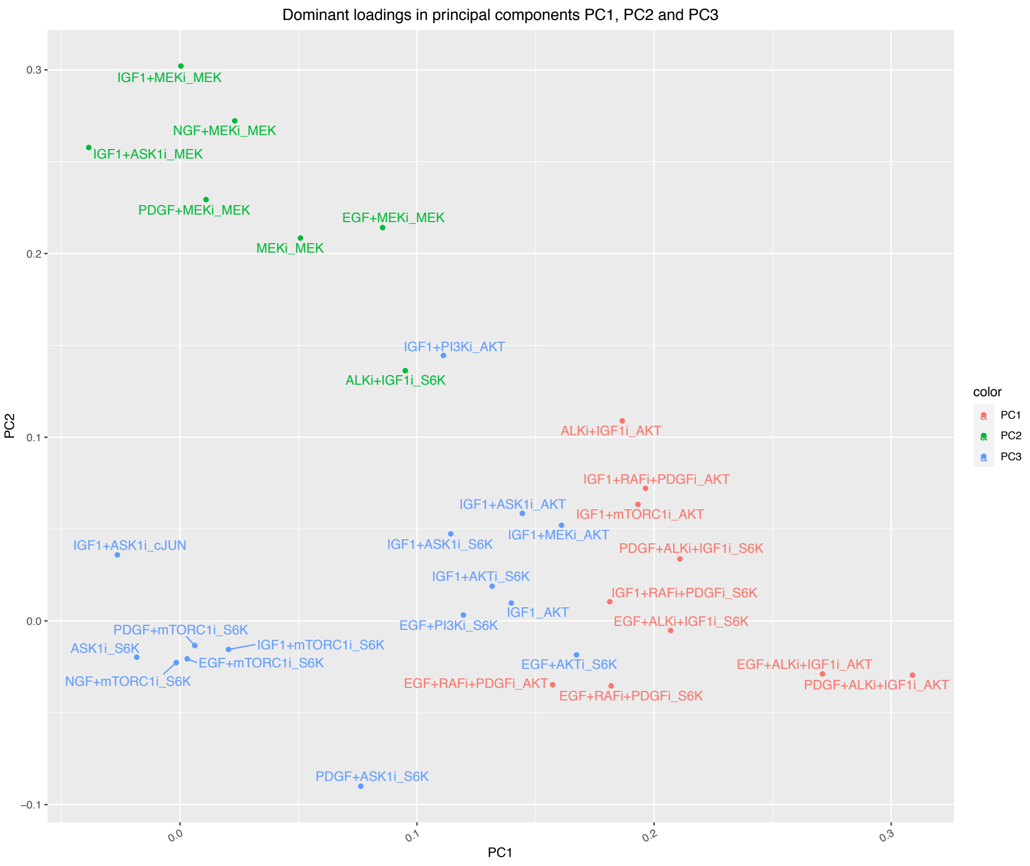


Supplementary Figure 10: Main loadings in the first 3 principal components of the perturbation data PCA. Color corresponds to the components for which the condition has the highest absolute weight.

| **Condition_PC1** | **PC1** | **Condition_PC2** | **PC2** | **Condition_PC3** | **PC3** |
|---|---|---|---|---|---|
| PDGF+ALKi+IGF1i_AKT | 0.309 | IGF1+MEKi_MEK | 0.302 | IGF1+ASK1i_MEK | -0.235 |
| EGF+ALKi+IGF1i_AKT | 0.271 | NGF+MEKi_MEK | 0.272 | IGF1+PI3Ki_AKT | -0.213 |
| PDGF+ALKi+IGF1i_S6K | 0.211 | IGF1+ASK1i_MEK | 0.258 | IGF1+mTORC1i_S6K | -0.210 |
| EGF+ALKi+IGF1i_S6K | 0.207 | PDGF+MEKi_MEK | 0.229 | PDGF+ASK1i_S6K | -0.209 |
| IGF1+RAFi+PDGFi_AKT | 0.196 | EGF+MEKi_MEK | 0.214 | PDGF+mTORC1i_S6K | -0.208 |
| IGF1+mTORC1i_AKT | 0.193 | MEKi_MEK | 0.208 | NGF+mTORC1i_S6K | -0.202 |
| ALKi+IGF1i_AKT | 0.187 | IGF1+PI3Ki_AKT | 0.145 | IGF1_AKT | -0.200 |
| EGF+RAFi+PDGFi_S6K | 0.182 | ALKi+IGF1i_S6K | 0.136 | EGF+PI3Ki_S6K | 0.183 |
| IGF1+RAFi+PDGFi_S6K | 0.181 | EGF+ASK1i_AKT | -0.118 | EGF+AKTi_S6K | 0.180 |
| EGF+AKTi_S6K | 0.167 | IGF1+MEKi_ERK | 0.117 | IGF1+MEKi_AKT | -0.179 |

Supplementary Table 1: Main loadings of the first 3 principal components of the perturbation data PCA.

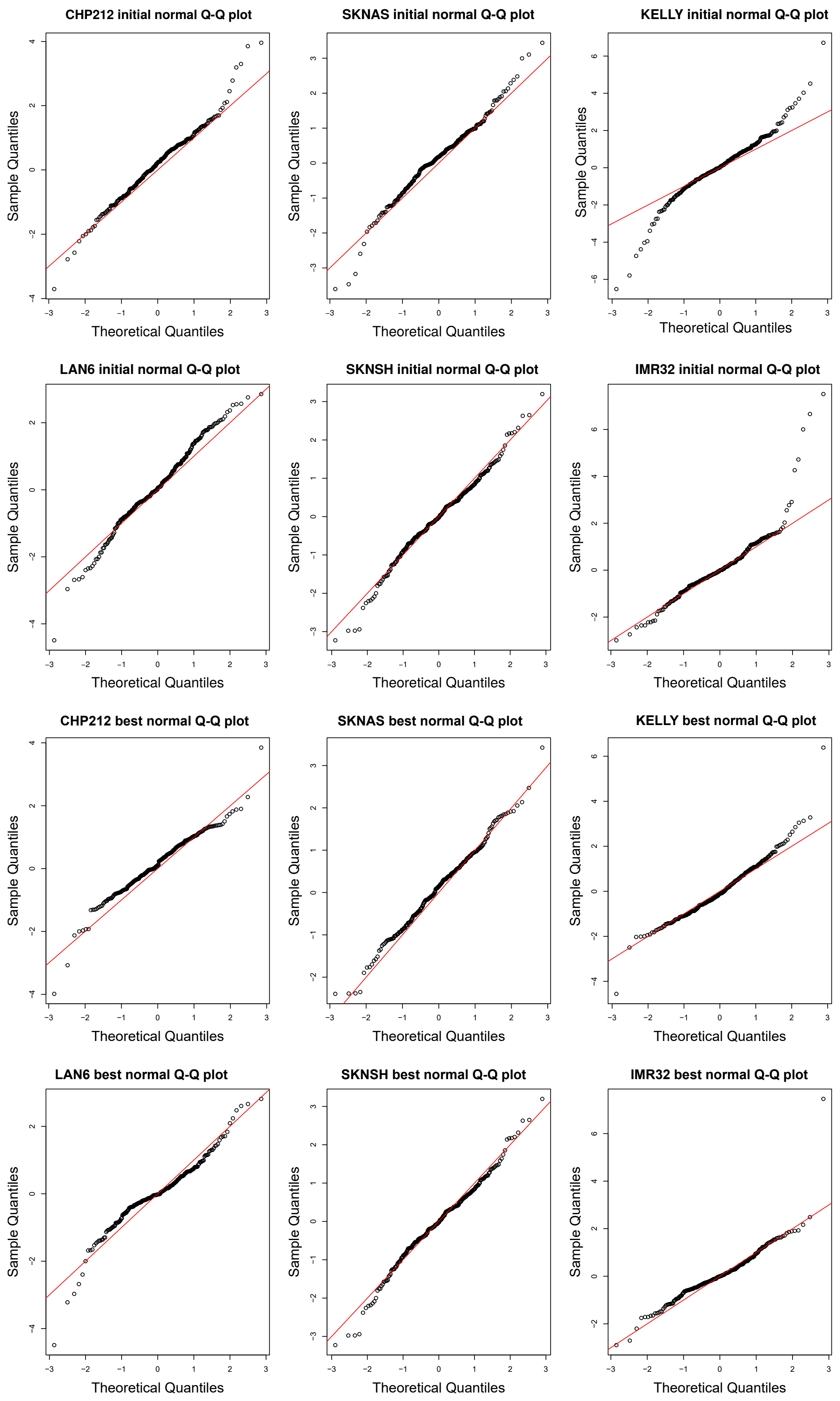


Supplementary Figure 11: Quantile-quantile plots of the initial models using the literature topology (top, initial normal)and the final models after model extension (bottom, best normal).

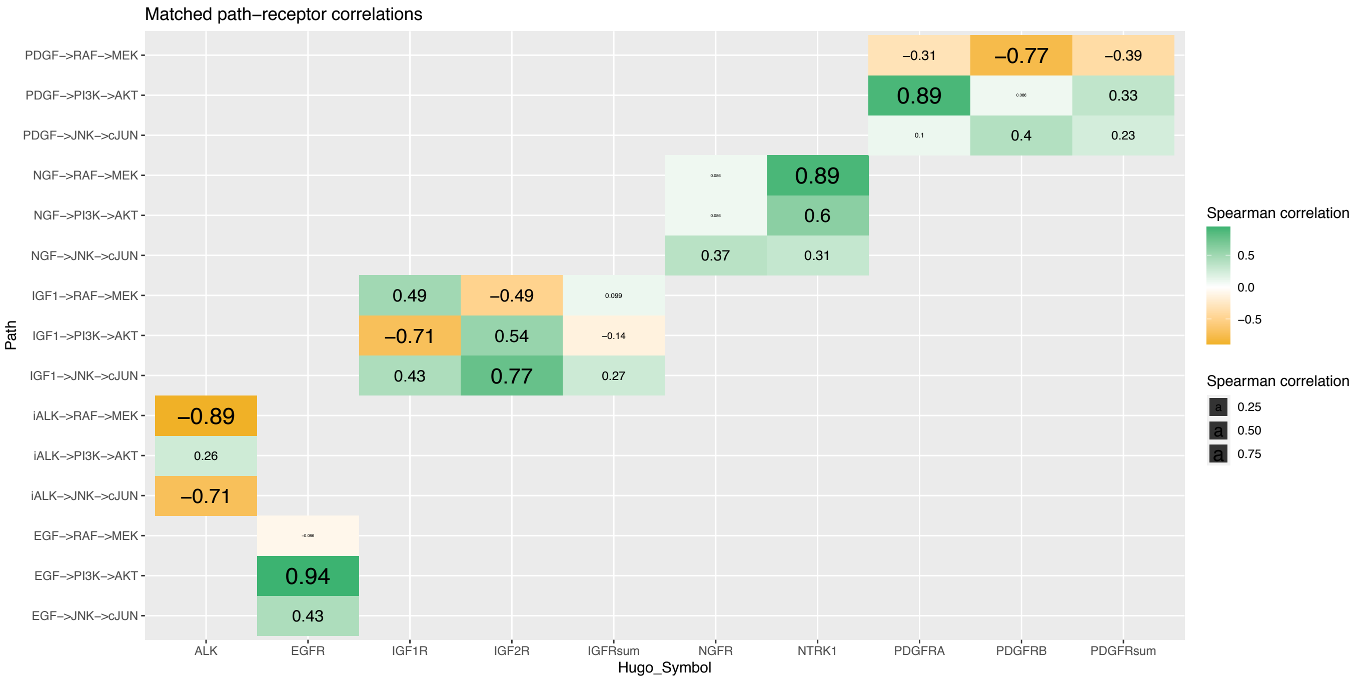


Supplementary Figure 12: Correlation between the fitted path value from ligands to readouts and the expression of the matching receptor or receptor family.

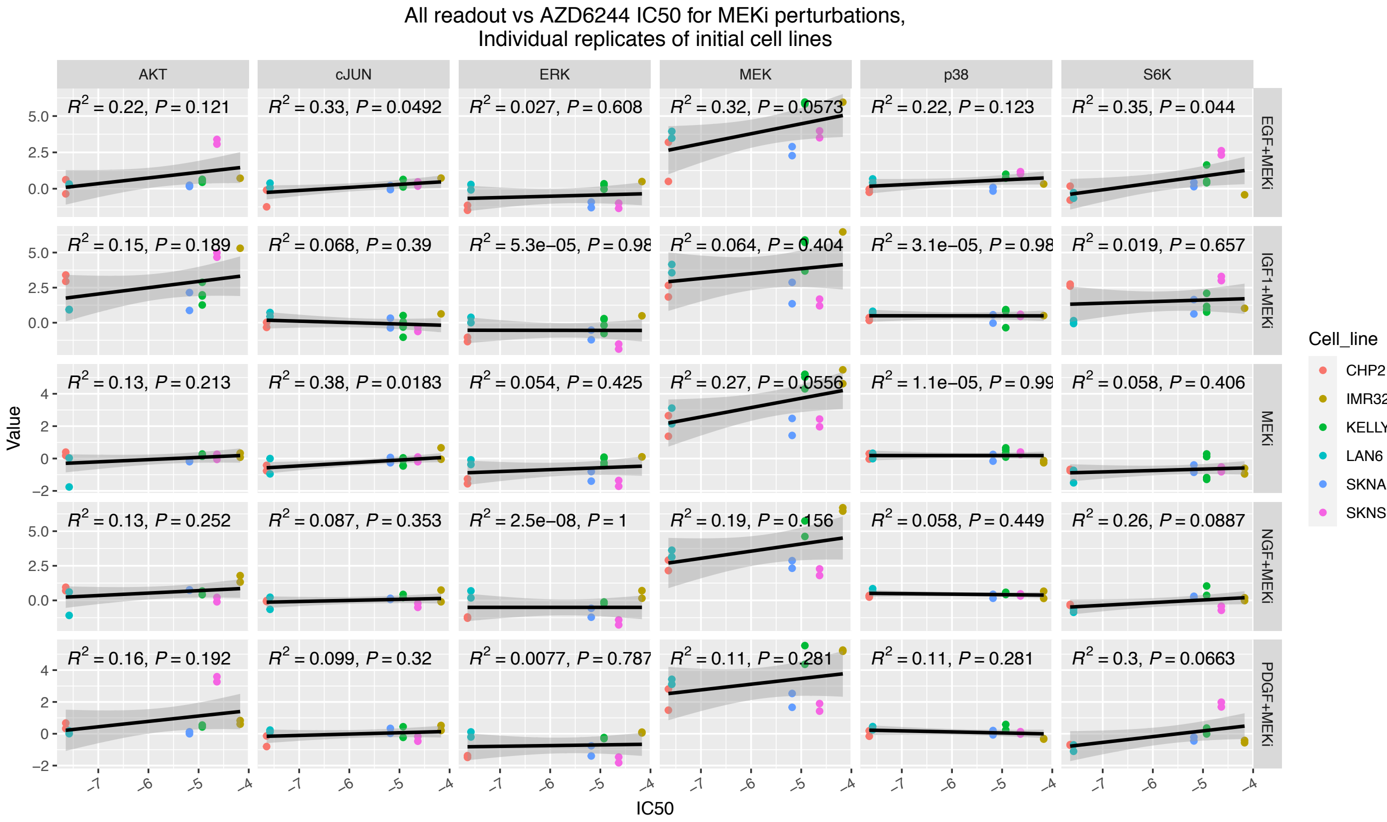


Supplementary Figure 13: AZD6244 IC50 versus responses to perturbations including AZD6244. $R^2$ and p.value correspond to the linear model shown in black. Points are independent replicates, n=2.

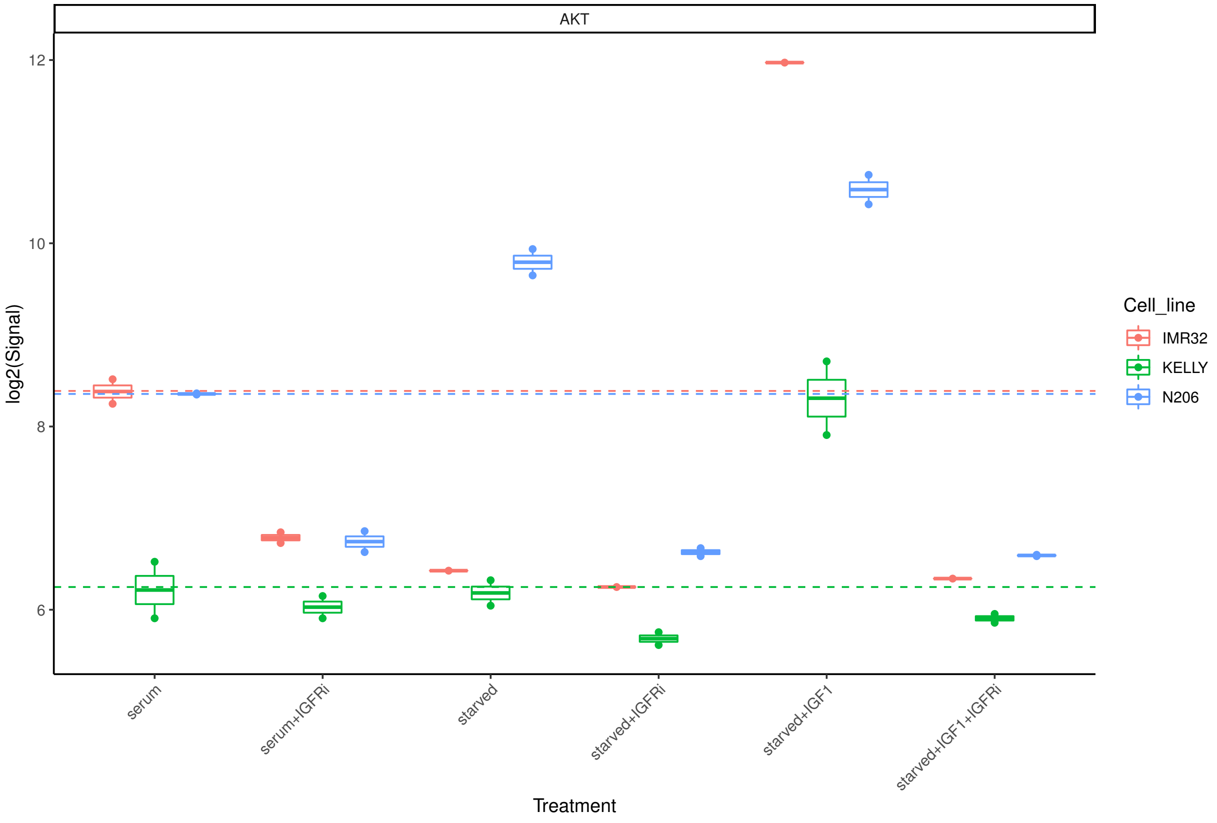


Supplementary Figure 14: Raw plex measurements of pAKT$^{S473}$ response to IGFR inhibition under serum, starved or IGF1 condition. The AKT pathway is activated by IGFR in N206 even without serum and is not activated more by IGF1 while IMR32 and KELLY requires IGF1 or other ligands. Dotted line indicates level under serum condition, n=2.

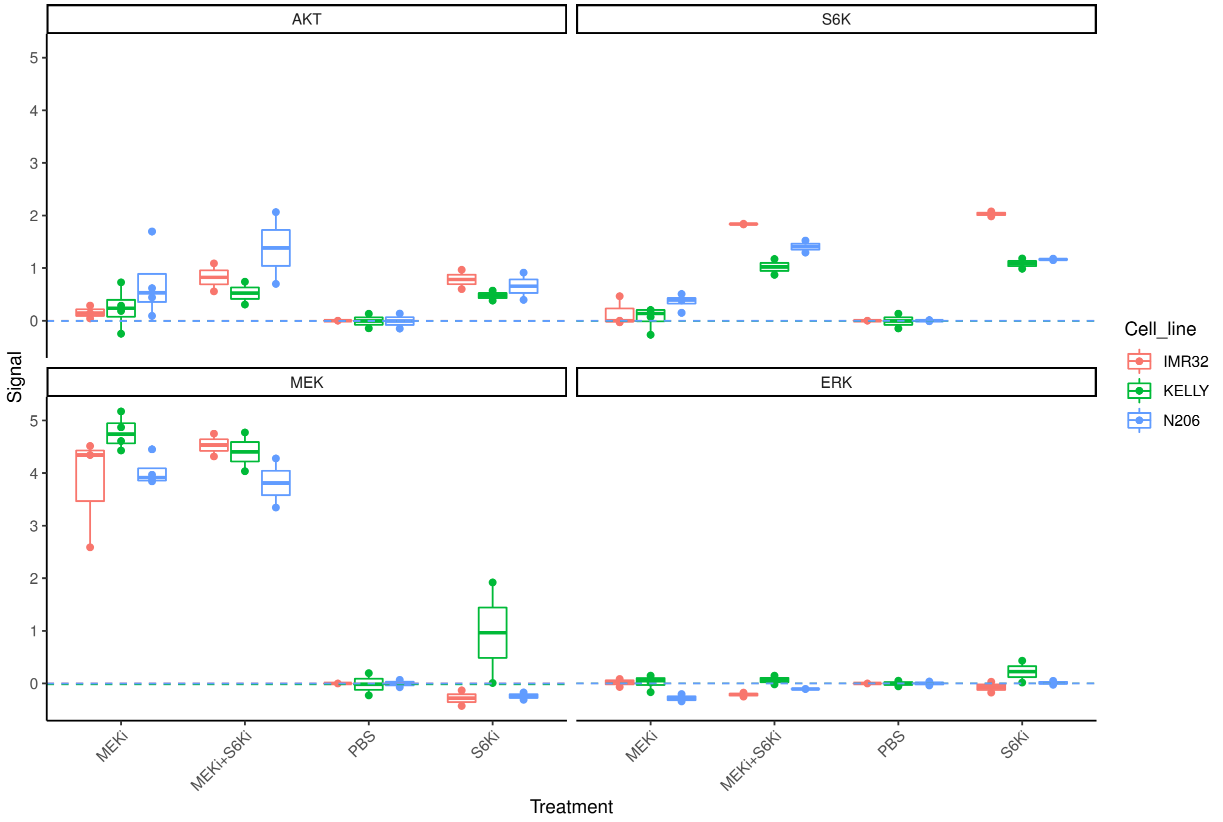


Supplementary Figure 15: PBS normalised plex measurements of p-AKT$^{S473}$, p-ERK$^{T202/Y204}$, p-S6K$^{T389}$ and p-MEK$^{S217/S221}$. n=2.

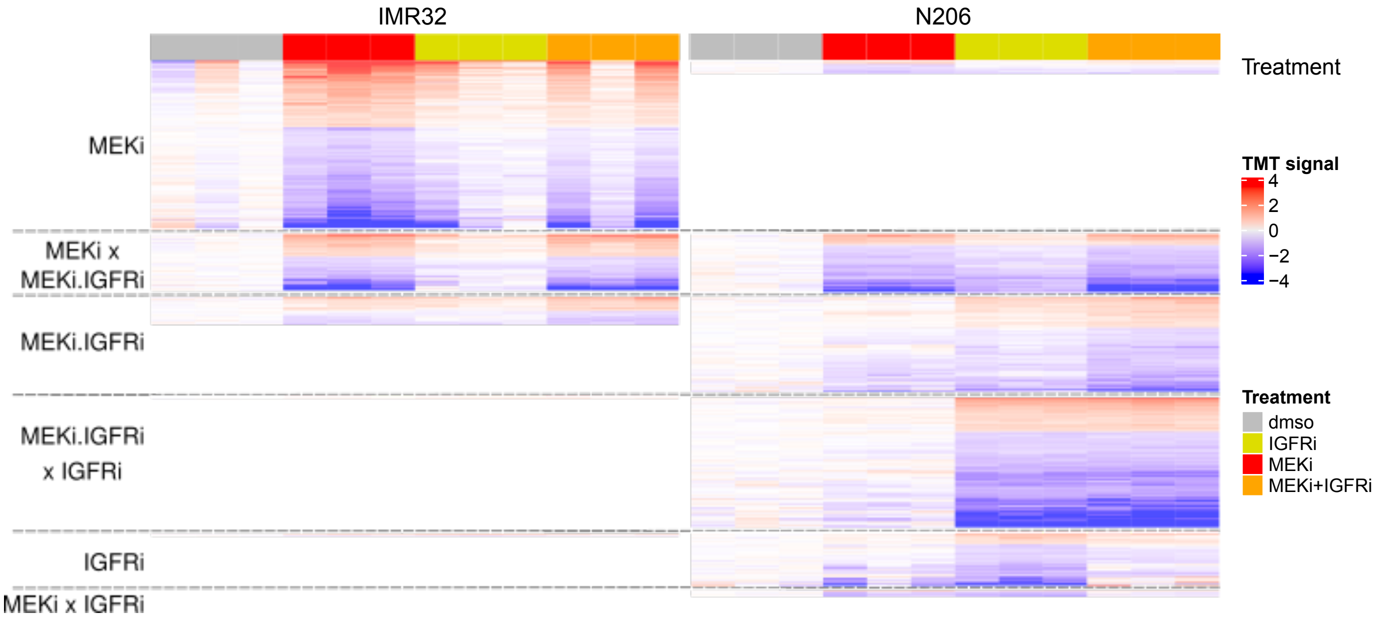


Supplementary Figure 16: Differentially measured phosphopeptides in IMR32 and N206 after 4h inhibition (FDR < 0.05, n=3) classified by the treatment(s) yielding a differential expression.

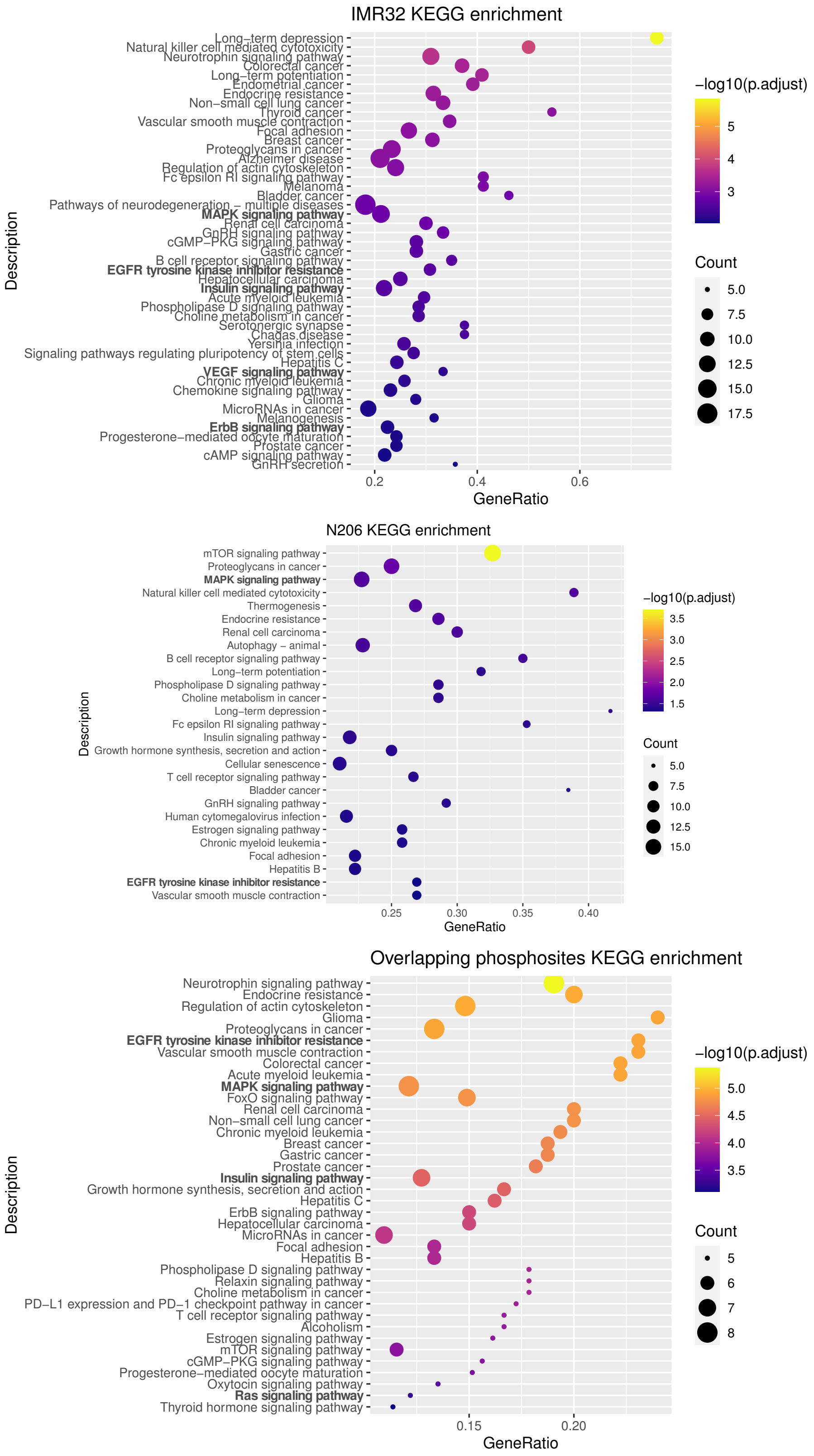


Supplementary Figure 17: KEGG enrichment of unique genes corresponding to phosphopeptides differentially expressed after MEKi, IGFRi or MEKi+IGFRi treatment in IMR32 (top), N206 (middle) or both strictly (bottom). MAPK related terms are highlighted. Computation using the R package enrichKEGG.
